# Sound-Driven Motion of Human Cochlear Structures Depends on Organ of Corti Integrity

**DOI:** 10.64898/2026.10.19.759683

**Authors:** Paul Secchia, Christopher I. McHugh, Nam Hyun Cho, Aleksandrs Zosuls, Andrew Tubelli, Sunil Puria, Hideko Heidi Nakajima

**Author notes:** Corresponding authors: Hideko Heidi Nakajima, Paul Secchia.

## Abstract

Optical coherence tomography (OCT) was used to image and measure the passive sound-driven displacements of cochlear structures *in situ* through the intact round window membrane of fresh human cadaveric specimens at 12 to 30 hours postmortem. We find that the anatomical integrity of the organ of Corti (OoC) affects cochlear structural motion. OoC with high integrity (erect with supporting cells) have small osseous spiral lamina (OSL) motion. Conversely, OoC with low integrity (flattened without supporting cells) have large OSL motion. For the ratio of motion between the basilar membrane (BM) and OSL (BM/OSL), OoC with high integrity has large BM/OSL motion while a cochlear partition without OoC has small BM/OSL motion. Furthermore, the high-integrity OoC has reticular lamina surface motion that is lower than the BM, whereas low-integrity OoC has similar reticular lamina and BM motion. This relationship between OoC anatomical integrity and cochlear structural motion will have consequential effects on sound transduction.

**Statement of Significance:** Acoustic vibrations are transformed into neural signals within the cochlea, the mammalian organ of hearing. Our understanding of cochlear mechanics has been limited to laboratory animals, whose cochleae have significant anatomical and functional differences from humans. To bridge this gap, we imaged and recorded the motion of cochlear structures in intact human cadaveric cochlear specimens without disturbing the interior of the cochlea. We determined how the anatomical integrity of the Organ of Corti (that contains structures for sound transduction) affects cochlear structural motion.

## Introduction

The cochlea, the organ of hearing, is responsible for converting acoustic vibrations into neural signals. Essential to this process is the *cochlear partition*, which bisects the fluid-filled cochlear channels and contains the sensory epithelium of the inner ear. The cochlear partition includes the osseous spiral lamina **(OSL)**, bridge, limbus, organ of Corti **(OoC)**, basilar membrane **(BM)**, reticular lamina **(RL)**, tectorial membrane **(TM)**, and spiral ligament **(SL)** (Fig. 1). The mechanical properties and anatomical architecture of structures within the cochlear partition dictate basic properties of hearing, such as passive sensitivity and sharpness of frequency tuning. Therefore, human passive mechanisms, which can differ from laboratory animals, are foundationally important for our understanding of hearing and effects of pathology.

**Figure 1:**
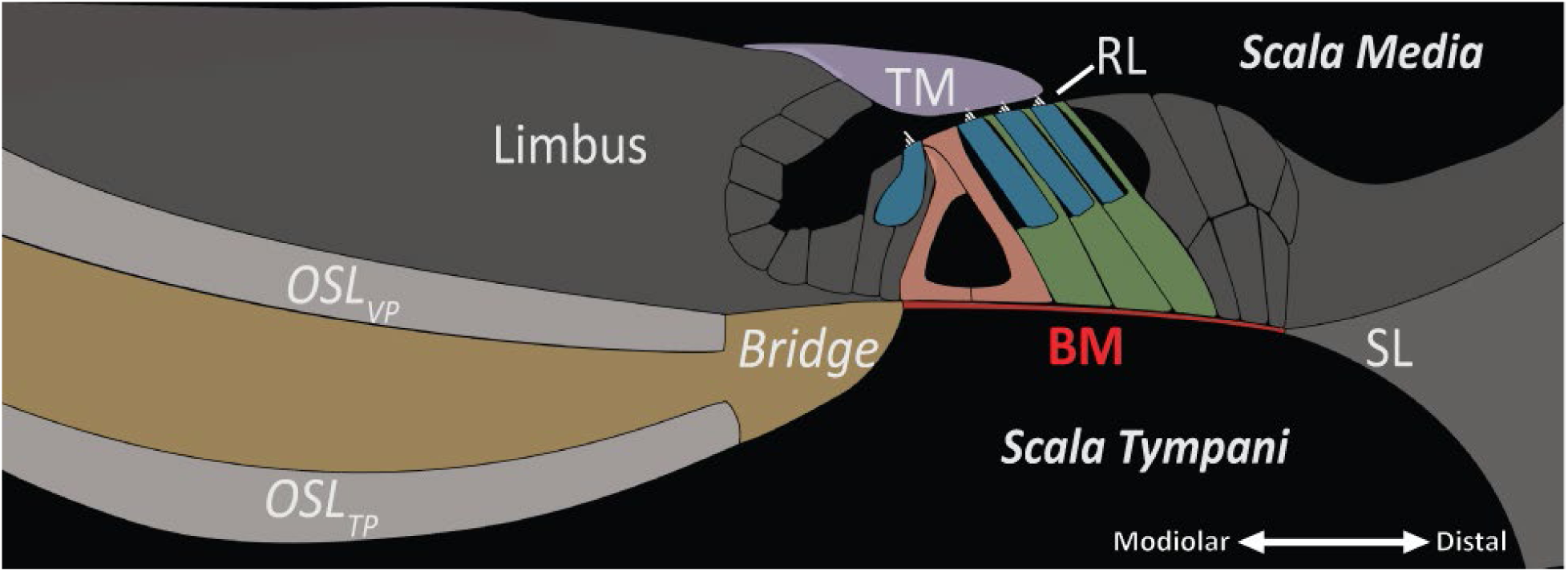
An illustration of the cochlear partition with labeled structures. Abbreviations: Osseous spiral lamina tympanic plate (OSL_TP_) and vestibular plate (OSL_VP_), basilar membrane (BM), tectorial membrane (TM), reticular lamina (RL), spiral ligament (SL).

The basal cochlear partition cross-sectional anatomy and structural motion were found to be similar across many species of laboratory animals (e.g. cat, chinchilla, rat, gerbil, and mouse). Thus, a *classic view* of cochlear mechanics emerged and was thought to be generalizable across mammals (1-3). In this classic view, the BM (with the OoC attached above) has been the mobile region of focus. In the classic view, the BM’s modiolar end attaches to the bony OSL and the BM’s distal end attaches to the spiral ligament. At these two ends of the BM, the OSL and the spiral ligament are considered rigid with negligible vibration, similar to other bony regions of the inner ear.

Although the classic view of mammalian basal cochlear mechanics was accepted for decades, deviations had been considered. Rhode (4) found that in squirrel monkey, the BM was not the only moving structure and that the OSL (beyond the BM) was flexible and moved. Although computational models of the classic view treated the OSL as an immobile and rigid structure (e.g., (5, 6)), there were notable exceptions. Allaire, Raynor, and Billone (7) predicted that due to the relatively long radial width of the OSL in human (10× the length of the BM in the base), relatively small transverse displacements of the OSL tip (in the direction perpendicular to the BM) could result in volume displacements of comparable magnitudes to that of the BM region. They concluded that the OSL would have a significant effect on the overall human cochlear partition stiffness in the basal region. Kohlloffel (8) also predicted that because the basal OSL is thin, fragile, and flexible in pig, cow, and humans, the basal OSL has the potential to deflect perhaps as much as the basal BM at low frequencies.

In cadaveric human, the OSL was reported as flexible from measurements made at two locations with laser Doppler vibrometry (9). More recently, Raufer et al. (10) made detailed laser Doppler vibrometry measurements of the scala tympani surface of the basal cochlear partition at multiple radial locations. They found that the thin wide tympanic plate of the OSL indeed bends with its distal end vibrating more than the modiolar end (like a plate anchored near the modiolus). Raufer et al. also noted anatomical features of the human cochlear partition that differed substantially from the classic view at the base (10, 11). Foremost among them is the presence of a soft-tissue “bridge*”* that connects the OSL to the BM (Fig. 1). In human, the limbus is attached to and straddles both the OSL and the bridge, and the limbus-TM attachment is located above the soft-tissue bridge. Thus, the TM-limbus attachment is likely not stationary as described in the classic view; rather, the motions of human cochlear partition structures are more complex.

To understand the intricate structural motions within the human cochlear partition, we measure the motion of various cochlear partition structures, normalizing them to the motion of the stapes. Normalizing to stapes motion instead of ear-canal pressure removes the variability introduced by middle-ear mechanics across ears as well as acoustic complexities of the ear-canal pressure measurements especially at high frequencies. With optical coherence tomography (**OCT**), we can both image and measure motion of specific locations of various structures *in situ* while keeping the cochlea and surrounding structures intact. A similar technique of measuring basal cochlear partition motion through the intact round window was reported by Kerkhofs et al. for frequencies below 5 kHz (12). Here, we focus our measurements on frequencies above 4 kHz, including the passive best frequencies (**BFpassive**) of the imaged regions of around 10-14 kHz.

## Methods

### Surgical Preparation

Fresh human cadaveric temporal bone specimens were obtained with consent at Mass General Hospital. Approval for this work was provided by Mass General Brigham (IRB #2020P000508, and protocols #2022P001102 and #2024B000033). Twenty seven temporal bones were used for this study. Ten previously-frozen bones were used to develop and refine experimental techniques for OCT vibrometry. Seventeen fresh human temporal bones were used for experimentation. Seven experiments were halted due to middle ear pathologies or trauma during specimen preparation. Cochlear structural motion at high frequencies (>10 kHz) were in the noise for five ears. For three of these five ears, high-frequency stapes motion were in the noise (possibly due to ossicular loosening (13, 14)). The other two ears without high-frequency cochlear motion had high postmortem times (>30 hours). This resulted in five remaining specimens (31-89 year-old donors, 12-30 hours postmortem) for the data presented in this manuscript.

Before collecting measurements, specimens were prepared similarly to previous studies (10, 15, 16). Temporal bones were extracted from the skull base as described in Nadol et al. (17). We then accessed the middle ear cavity by widely opening the facial recess while taking care not to disturb the ossicles or suspensory ligaments by touching with instruments or allowing a suction tip in the vicinity. The bony overhang surrounding the round window membrane (**RWM**) was reduced to maximize the visibility of the cochlear partition. Specimens were then affixed to an articulating and lockable holder (Noga, DG0036, Israel) coupled to a 3-axis linear translation stage to orient the bone for imaging. To preserve the specimen, the sound isolation chamber was cooled to approximately 18° C. The middle-ear cavity was kept moist with cooled saline-saturated gelatin sponge pieces (Gelfoam, Pfizer, USA) placed in regions of the middle-ear cavity in a manner that did not obscure the imaging view, affect middle-ear motion, or result in liquid pooling in the middle ear.

### OCT Imaging and Vibrometry

OCT measurements began within 12-30 hours after the death of the donor. A Spectral Domain OCT system (GAN620C1, Thorlabs, Germany) with 900-nm center wavelength, an A line-scan camera frame rate of 46 kHz, and 2.2 µm axial resolution (in water) and ∼8 µm lateral resolution (36 mm, 0.055NA, 2x objective lens) was used. Custom LABVIEW-based software (VibOCT v 2.1.5 – 2.1.9) recorded images and made vibration measurements along A-lines (18). A custom Labview-based data acquisition software program (SyncAv v 0.47) performed synchronous averaging on a National Instruments PXI data acquisition computer with AD/DA hardware. SyncAv also generated pulses to synchronize the VibOCT camera with zero crossings of the tone generator. Pure-tone sequences from 0.1-21 kHz equalized for near constant stimulus levels, ranging from 70 to 120 dB SPL, were presented with a speaker (Radioshack 40-1377) in the open ear canal. Ear-canal sound pressure was measured with a probe-tube microphone (Knowles EK23103) with the probe-tube tip approximately 1-2 mm from the center of the tympanic membrane.

After mounting the specimen on the lockable holder, we visualized the cochlear partition through the intact round-window membrane with OCT. The specimen was oriented to generate a cross-sectional image of the cochlear partition (as depicted in Fig. 2), with the optical axis aligned approximately normal to the BM. This was done to collect vibrometry measurements made at 1024 points along A-lines in response to sound stimuli presented in the ear canal. This was repeated for 20-26 different radial locations along the cochlear partition. Reported vibrometry data of cochlear structures were at least 9 dB above the measured noise floor for each frequency, resulting in reliable displacement measurements above 0.1-0.4 nm. Noise-floor/artifact measurements of probe-tube microphone ear-canal pressure and OCT cochlear partition displacement were collected with the same set-up but with almost no electrical drive to the speaker (∼2×10^-15^ fV), thus any artifactual electrical pick-up throughout the set-up would be understood. We also ruled out motion artifacts of the specimen by verifying that the OCT cochlear-promontory displacement was at the noise floor when loud stimuli (70-120 dB SPL) were presented within the ear canal. Vibrometry data presented in this manuscript are above 4 kHz, where we confirmed that artifacts from round window motion on OCT measurements were negligible.

**Fig. 2.**
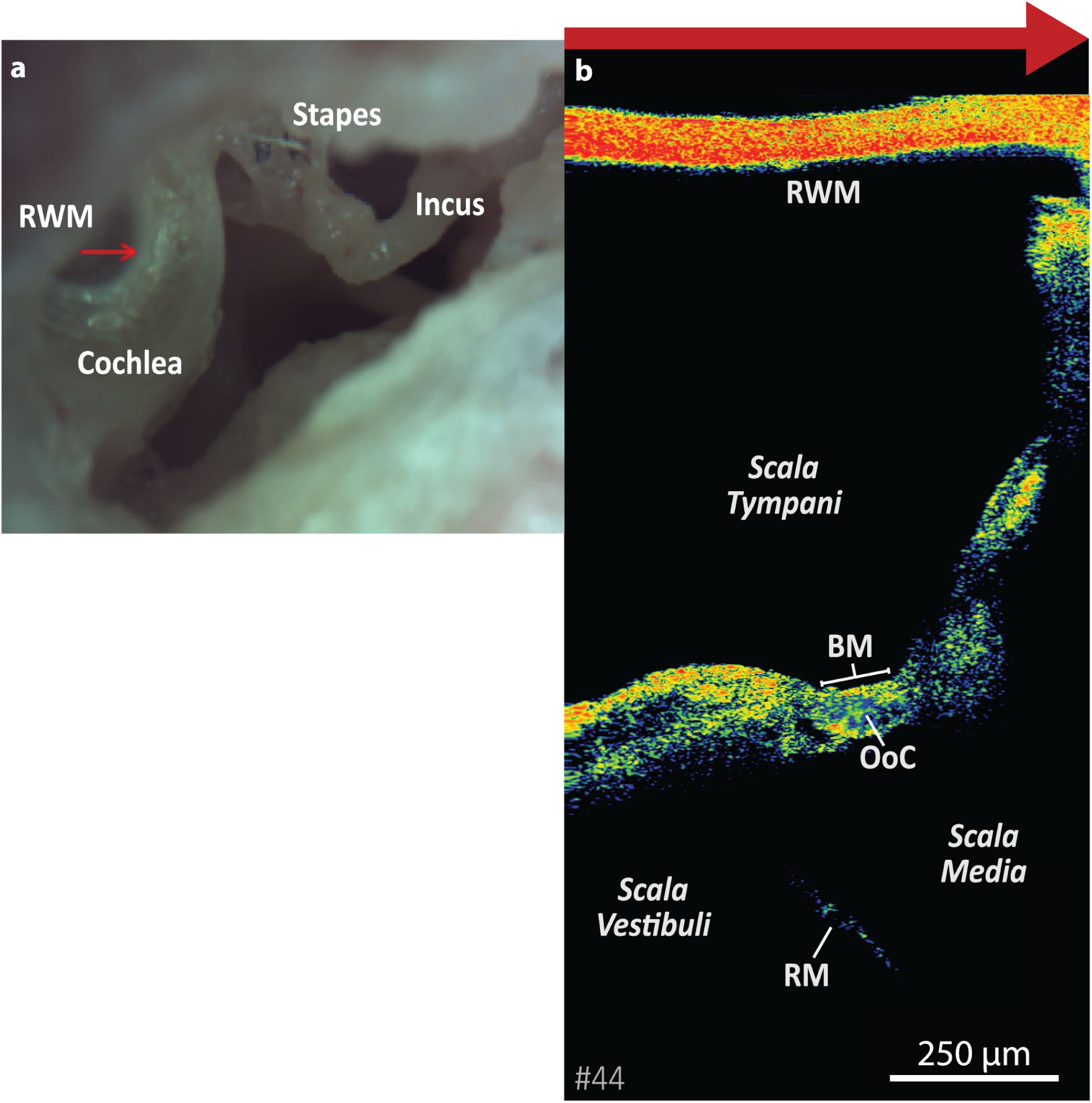
(a) Microscope view of the middle-ear cavity with a view of the round window and middle ear ossicles. The red arrow at the round window membrane shows the length and direction of the OCT scanning path. (b) OCT image of Ear A with corresponding red arrow at the top. Below is the round window membrane (RWM), the fluid filled scala tympani, and the cochlear partition with basilar membrane (BM) and organ of Corti (OoC). RM=Reissner’s membrane

Stapes velocity was measured with a laser Doppler vibrometer (CLV 700, Polytec, Germany), and after converting to displacement, was used to normalize the OCT displacement measurements of cochlear structures. Stapes displacement measurements were confirmed to be linear for the sound levels (less than 120 dB SPL) and frequencies used. The magnitudes of the frequency responses for stapes displacement were within normal limits (14).

## Results

### Visualizing the Cochlear Partition with OCT

Figure 2a shows the OCT camera view of the middle-ear cavity. The bony cochlea and parts of the middle ear ossicles and RWM are visible. The red arrow over the intact RWM is an example scanning path of the OCT beam. It indicates one dimension of the OCT two dimensional B-scan, where the second dimension is through the RWM and into the cochlea. Figure 2b shows the OCT B-scan image with the large red arrow at the top that corresponds to the red arrow of Fig. 2a. In Fig. 2b, the OCT light beam is directed from top to bottom and scans along the direction of the red arrow (left to right) to obtain the B-scan image. The ear in the OCT image is referred to as Ear A.

In previous work, we estimated the location of the OCT-imaged region of Ear A to be 3.56 mm from the base by histologically processing and graphically reconstructing the contralateral ear from the same donor (16). This corresponded to an estimated BFactive of 12.3 kHz when calculated with Greenwood’s cochlear tuning map for human (19). For the other ears in this study, we estimated that we imaged 1-5 mm from the very base, which corresponded to the ∼9.5-20 kHz BFactive region in a live active cochlea (which is half an octave above the cadaveric BFpassive).

Figure 3a shows illustrated estimations of anatomical structures superimposed over an OCT B-scan image. Figure 3b shows the OCT image without the illustration. This OCT image was collected 13 hours postmortem from Ear A. (The images in Fig. 3a&b are flipped compared to Fig. 2b, thus the OCT light passes from the bottom to top.) The bright orange and yellow colors indicate high OCT light reflectivity from tissue such as bone of the OSL, and as a result the OSL shadowed much of the limbus that was above it. The OSL-bridge junction (**OSL-Bridge-J**) is marked with a green dashed line in Fig. 3b, and can be identified by the border between the OSL and where the OCT light can penetrate through the soft-tissue bridge (low reflectivity and lack of shadowing artifacts into the limbus). The bridge-BM junction (**Bridge-BM-J**), marked with solid orange line in Fig. 3b, was identifiable due to an angle between the intersection of bridge and BM.

**Fig. 3.**
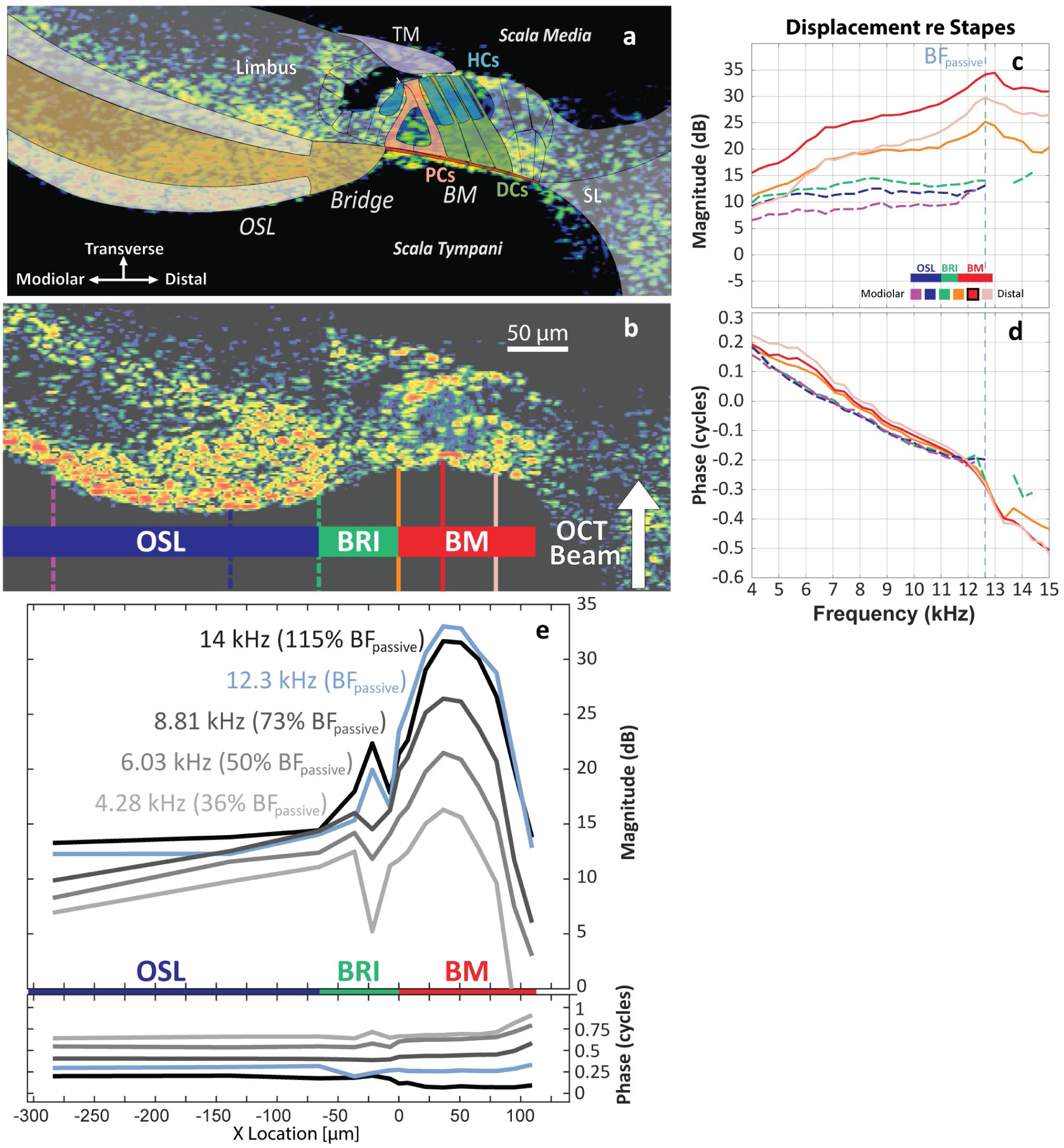
Example of OCT B-scan image and displacement data for Ear A: (a) Illustration detailing cochlear structures of interest over OCT image. (b) OCT B-scan image with radial locations of measurements. The OCT beam is projecting through the intact round window from below onto and through the tympanic surface of the partition. (c-d) Displacement of the tympanic surface of the cochlear partition normalized to stapes displacement is plotted for the magnitude and phase against frequency. Different radial locations are indicated with colored lines that correspond to the vertical lines in (b). The peak in magnitude and rapid phase change in (c-d) indicate that the passive best frequency (BF_passive_) was near 12.3 kHz (vertical blue dashed line). (e) Displacement of OSL, Bridge, and BM normalized to stapes displacement (Y-axis) plotted as a function of radial location (X-axis) where the Bridge-BM-J is set to X=0. Blue lines represent motion at BF_passive_, black lines are above BF_passive_, and different shades of gray are frequencies below BF_passive_. All data plotted have >9dB SNR.

### Motion of the tympanic surface of the cochlear partition

Sound-induced motion of various cochlear structures was captured with OCT. All motion measurements were approximately in the transverse direction (direction indicated in Fig. 3a). Figure 3b, with colored vertical lines, describes example radial locations along the cochlear partition’s tympanic surface (bottom surface in Fig. 3b) where motion was analyzed. The tympanic surface displacements normalized to stapes displacement are plotted as a function of frequency for the magnitude (Fig. 3c) and phase (Fig. 3d). The different colored lines correspond to the different radial locations demarcated by the matching-colored vertical lines in Fig. 3b. The largest transverse displacement across radial locations was seen at the BM (red solid line, Fig. 3 b&c). Other radial locations along the scala tympani surface, such as along the OSL (green, blue, and purple dashed lines), had lower magnitudes, in accordance with previous LDV measurements of the basal cochlear partition surface (10) and model calculations (20).

The BFpassive from each experiment was determined from the displacement frequency response where there was a peak in the magnitude and corresponding phase transition. For example, the ear in Fig. 3 c&d had a peak BM displacement and sharp drop in phase near the BFpassive of 12-13 kHz (marked by the dashed light blue vertical line). In ears without a clear peak in magnitude or phase transition, the BFpassive was estimated as the highest frequency where displacement continued to increase (with >9dB SNR), and where there were no irregularities in measurements of ear canal pressure and stapes displacement. The reported measurements all came from ears that were confirmed to have robust stapes motion at high frequencies. The BFpassive for all specimens was between 11-13 kHz.

Figure 3e shows the displacement magnitude and phase of the tympanic surface of Ear A as a function of radial location (X-axis) at several frequencies (different colored lines). The radial location X=0 µm corresponds to the Bridge-BM-J (solid orange line in 3b), thus points with a positive X-value are located in the BM and points with negative X-value are located in the OSL and bridge. The motion of the BM increases with frequency when approaching BFpassive, with a peak displacement of about 33 dB normalized to stapes displacement. The location of maximum displacement was near the estimated location of the outer pillar cell foot, and this location also had the maximum displacement at frequencies below and above BFpassive.

Generally, the OSL had greater displacement magnitudes at higher frequencies. At the bridge (near X=−25 µm), there was a localized increase in motion with some phase change at BFpassive=12.3 kHz and at 14 kHz. But at lower frequencies, this bridge location had a localized decrease in motion.

OCT B-scan images of 5 representative specimens (Ears A-E) are shown in Fig. 4 in order of high to low anatomical integrity (16). There was substantial variation in the OoC anatomy across these ears. Ears A and B had large and erect OoCs where the surface of the TM closely interfaced the RL. Ear C also had an erect OoC, but with a prominent void within the OoC that may be due to hair cell loss (16). Ear D had a flattened OoC, which indicates a lack of supporting cells, as described in previous histologic studies (16, 21, 22). Also, the TM was lifted off from the OoC structures in Ear D. Ear E lacked an OoC altogether. Figure 5a plots displacements of the tympanic surface of the cochlear partition normalized to stapes displacement (Y-axis) against radial location normalized with respect to the Bridge-BM-J (X-axis) at BFpassive for Ears A-E. These plots are similar to Fig. 3e where radial location of X=0 corresponds to the Bridge-BM-J and is marked by the vertical dashed black line. To the left of the Bridge-BM-J (located at X=0 µm), motion is plotted for the bridge and even further to the left, the OSL.

**Fig. 4.**
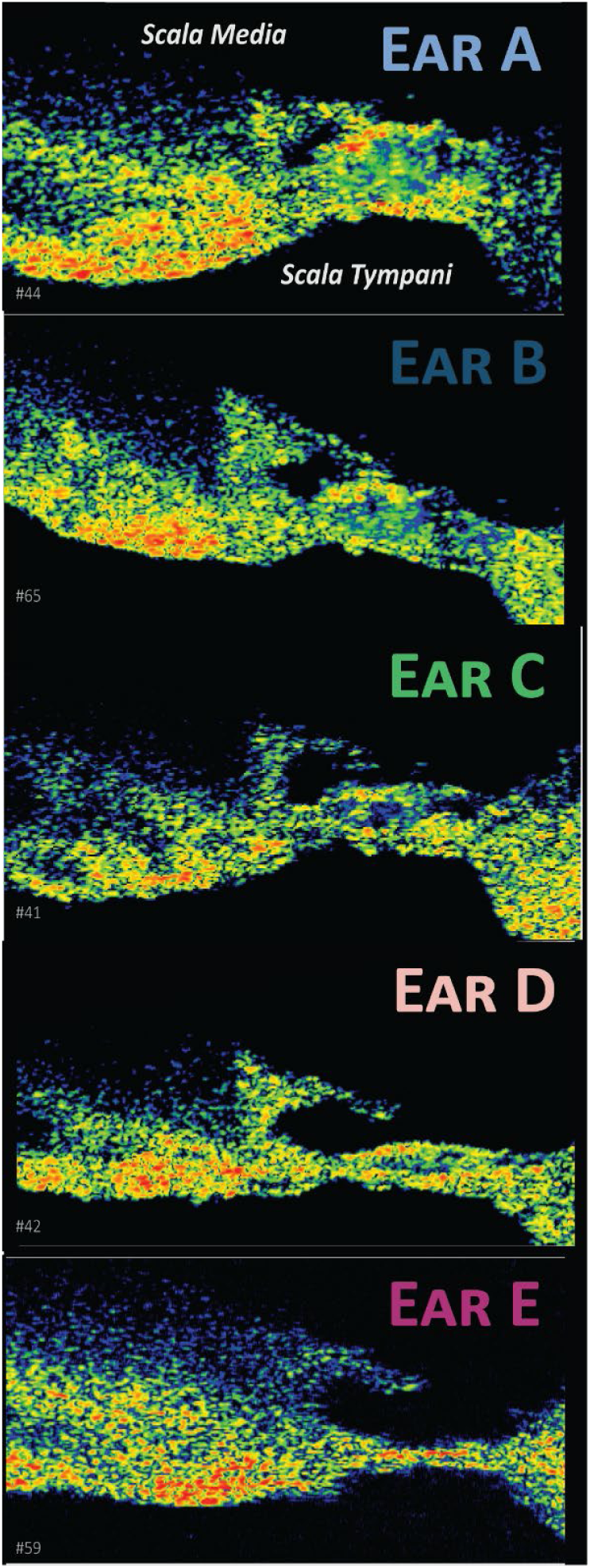
OCT B-scan images. Images of each ear are arranged in order of largest to smallest organ of Corti (OoC) size. Ears A and B are considered to have high cochlear integrity due to the normal, erect shape of the OoC. Ear C also has an erect and normally shaped OoC, however it contains a prominent empty space within the OoC (likely due to lack of hair cells). Ear D has a flattened OoC while Ear E has no visible OoC, and thus Ears D-E are of low cochlear integrity and lack supporting cells (16). In Ears A-C the TM surface interfaces closely with the reticular lamina (RL), while in Ear D the TM is lifted away from the RL.

**Fig. 5.**
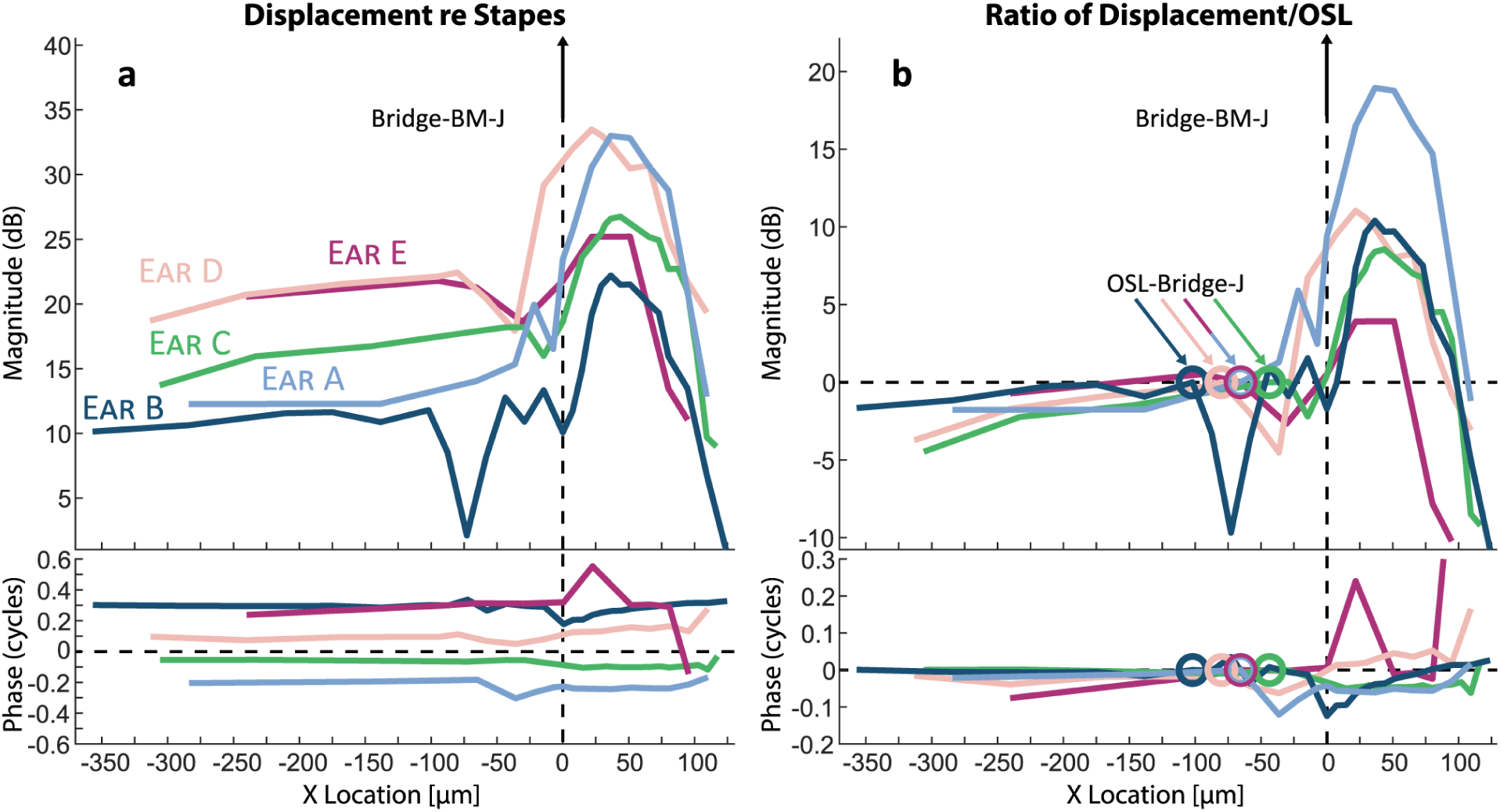
(a) Transverse displacement at BF_passive_ along the tympanic surface of the cochlear partition normalized by the stapes displacement. This displacement (Y-axis) is plotted against different radial locations where the X-axis is the referenced distance with respect to the Bridge-BM Junction (Bridge-BM-J). Colored lines represent different ears with B-scan images shown in Fig. 3. (b) Displacement ratio of structures/OSL plotted versus radial location. The “OSL references” are the displacements at the OSL-Bridge Junction (OSL-Bridge-J) (circles at Y=0). (Note that only about 240-350 µm of the 1300 µm total OSL length was visible through the round window in these experiments.)

Fig. 5a shows that overall, the radial profile of motion was similar across ears. However, Ear D slightly differed from the other ears where the radial motion profile appears shifted to the left, and the peak displacement is closer to the Bridge-BM-J. The OSL (to the left of the bridge, X<−100 µm) was mobile and its motion normalized to stapes displacement varied across ears between 10-22 dB. Interestingly, cochlear partitions with larger and fuller OoCs (Ears A & B plotted with light and dark blue) exhibited the lowest OSL displacements (10-12 dB) as compared to other ears that appeared to have had some pathology. Ear D with flattened OoC and Ear E with no OoC had the highest OSL motion (∼20-22 dB), while Ear C (which had an erect OoC containing empty spaces) had an OSL motion in-between (∼13-18 dB). The OSL moved less in ears with greater anatomical integrity. In all ears, the point of maximum motion was at the BM near the estimated location of the foot of the outer pillar cell. The maximum displacement of BM motion normalized to the stapes ranged between 22-34 dB. The phase remained smooth and similar across radial locations of the OSL and BM. However, there were phase changes at different radial locations throughout the soft-tissue bridge region.

Fig. 5b plots the displacement ratio of each measured location referenced to the displacement at the OSL-Bridge-J. The OSL-Bridge-J is marked by open circles for each ear, and whose motion was set to 0 dB in the Y-axis. (Note, the radial dimensions of the BM and OSL widths vary across ears). Points to the right of the Bridge-BM-J (X>0 µm) are located within the BM, and their magnitude of motion is a ratio of “BM/OSL” displacement. In Fig. 5b, one can appreciate the large variations in BM/OSL displacement ratio across ears. In Ear A (which had the largest OoC), the maximum BM/OSL ratio was 19 dB. Ears B, C, D had BM/OSL ratios of 8-11 dB, whereas Ear E (with no OoC) had a BM/OSL of only 4 dB. It appears that the ear with the best cochlear integrity (Ear A) had the highest BM/OSL displacement ratio, while the ear with the worst integrity (Ear E) had the lowest BM/OSL ratio. Ears B, C, D had similar BM/OSL ratio magnitudes. The BM/OSL phases were negative for Ears A, B, C (ears with good cochlear integrity), whereas the BM/OSL phases were more positive for Ears D and E (ears with poor cochlear integrity).

### Motion of cochlear partition structures

One of the advantages of OCT is that it can measure motion of cochlear structures beyond and through the tympanic surface of the cochlear partition, such as the RL and TM. Figure 6a is an OCT image of Ear A with multiple markers designating the selected locations where vibrometry measurements were analyzed. Only a few locations within each A-line are plotted for simplicity, and to remove potential bias, these locations were selected by an automated algorithm that identified points of high signal-to-noise motion near the surfaces of the cochlear partition. Different structures are designated by marker shape and color. The square markers are along the tympanic surface of the cochlear partition and consist of the OSL (blue), bridge (green), and BM (red). The orange triangles denote measurement points along the RL, whereas the purple circles are points in the TM.

**Fig. 6.**
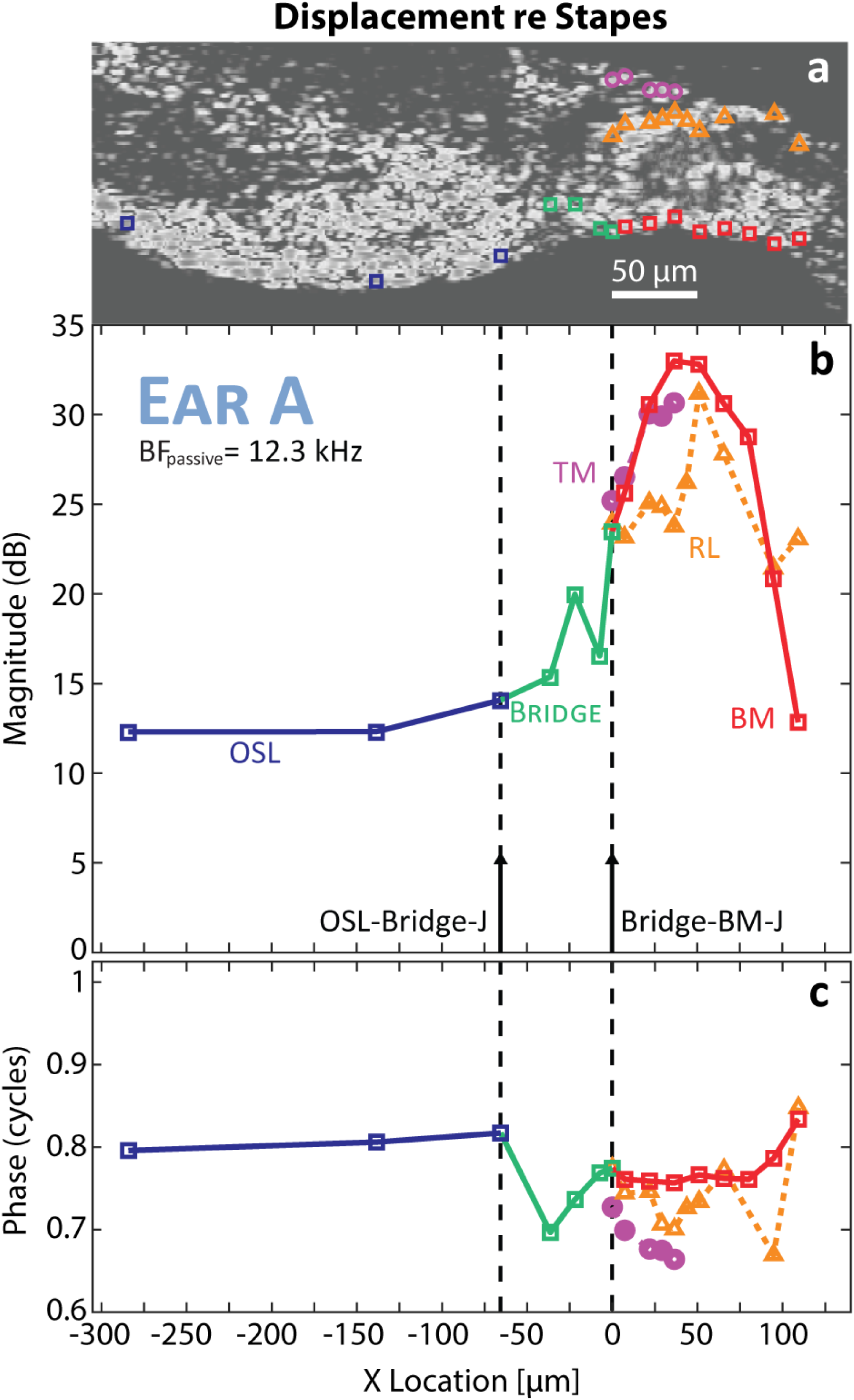
Motion of cochlear structures vs radial position at BF_passive_: (a) OCT B-scan image and location of vibrometry measurement points (color and shape-coded by structural regions). Vibrometry data of these points are plotted in b-c. Displacement magnitude (b) and phase (c) normalized to stapes displacement are plotted against radial position for Ear A. A radial location of X=0 corresponds to the Bridge-BM-J.

Figures 6b-c plots magnitude and phase of point transverse displacements normalized by stapes displacement of cochlear structures plotted against radial location at BFpassive for Ear A. A radial location of 0 µm corresponds to the Bridge-BM-J, indicated by the vertical dashed line on the right. A second vertical dashed line on the left indicates the OSL-Bridge-J, which was about 65 µm away from the Bridge-BM-J. The magnitudes of the OSL displacement (blue squares) were approximately 12 dB. The motion of the bridge increased to about 23 dB at the Bridge-BM-J. The BM displacement (red squares) reached a peak of about 33 dB near the foot of the outer-pillar cell. Distal from this peak towards the spiral ligament, the BM motion decreased to about 12 dB. The motion of the RL (orange triangles) reached its maximum displacement at X=50 µm. The displacement magnitude of the RL varied from 2-10 dB lower than the BM. The magnitude of TM motion (purple circles) was similar to the BM (<2.5 dB difference). The phase of the BM was smooth with respect to radial location. However, the phase of the RL and of the bridge fluctuated with location.

Figure 7 contains OCT images and vibrometry displacement plots for Ears B-E (similar plots to Fig. 6). The maximum motion of the OSL was at its distal edge (the OSL-Bridge-J) for all ears, consistent with Raufer et al. (10), and as expected. The magnitude and phase of OSL motion was similar at the tympanic plate (dark blue squares) and at deeper locations within the OSL (near the vestibular plate, plotted with lighter blue triangles and circles). The bridge (green squares and yellow triangles) had more variation in its magnitude and phase across location and across ears, and seemed to act as a transition region between the OSL and BM. The maximum displacement of the cochlear partition re stapes ranged between 22-33 dB and was within the BM near the location of the foot of the outer pillar cells.

**Fig. 7.**
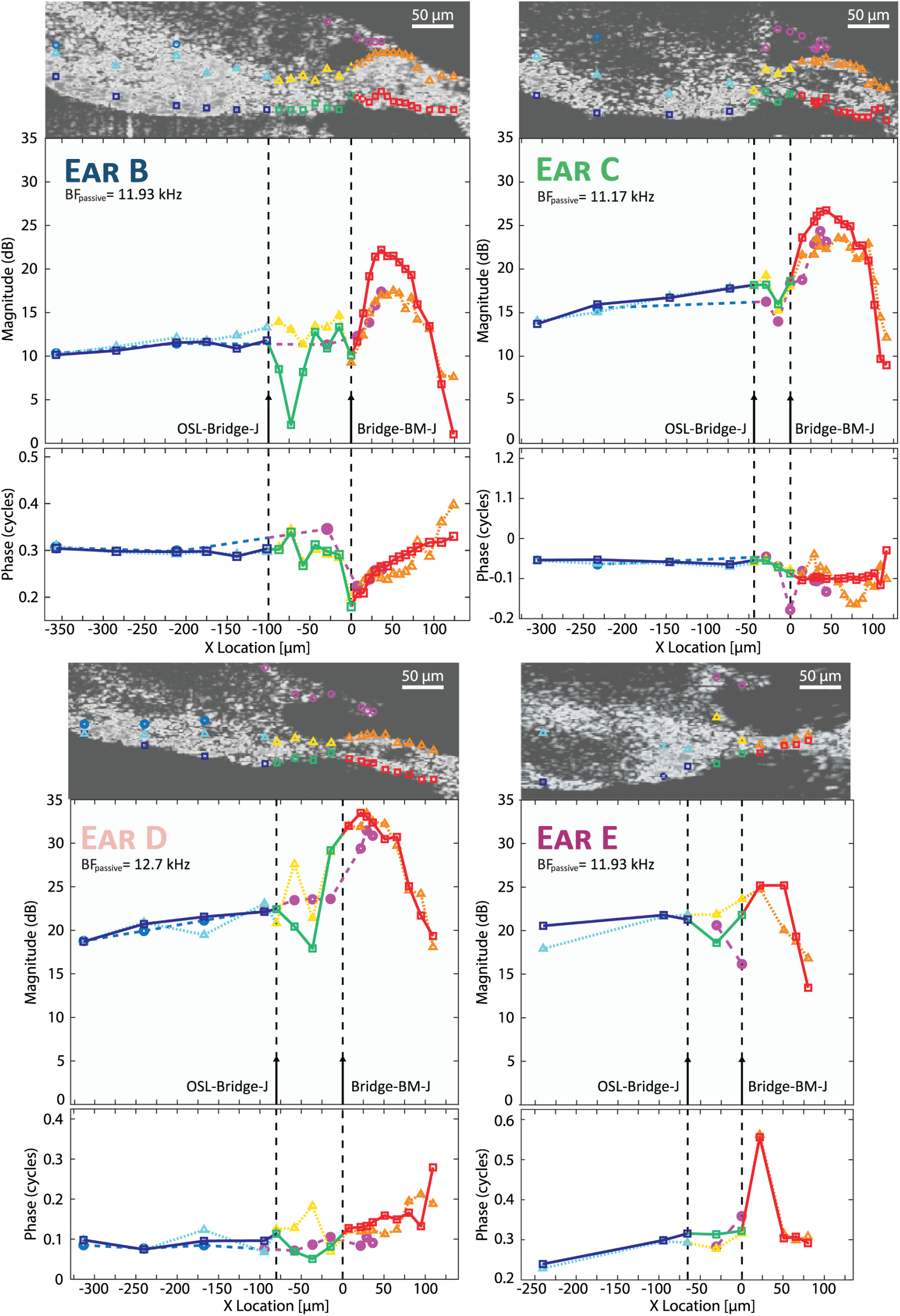
Displacement of cochlear structures vs radial position at BF_passive_ for Ears B, C, D, and E. See Fig. 6 for details.

In Ears A-C (with erect OoC), the motion of the RL (orange triangles) was approximately 4-10 dB less than the BM. The RL moved about the same magnitude as the BM in Ears D-E (flat or no OoC). However, the RL phase fluctuated while the BM phase was smooth across radial locations. In Ears B and C, the TM displacement was similar to the RL, with similar magnitude and fluctuating phase with location.

## Discussion

In fresh human specimens 12-30 hours postmortem, motion of cochlear partition structures is measured *in situ* by visualizing through the intact RWM without opening the cochlea or inserting instruments into it. OCT allows simultaneous imaging and vibrometry of cochlear structures, allowing measurements of sound evoked motion of specific anatomical structures. Furthermore, we ensure our analyses were based on displacement measurements with good signal-to-noise (>9 dB SNR) and free of artifacts. We avoid analyzing data of frequencies below 4 kHz where the magnitude of motion of the RWM is larger than the cochlear partition by more than about an order of magnitude (23). Since the OCT beam travels through air above the RWM and through fluid below the RWM, RWM motion introduces a moving air-fluid interface that alters the optical path length. This artifact dominates motion measurements made at the base at low frequencies, where RWM motion is greatest and cochlear partition motion is small (24, 25).

Previously, we reported that variations in cochlear anatomy and anatomical integrity of the OoC can be estimated by OCT imaging through the intact RWM (16). Key among the factors for determining anatomical integrity was the size and shape of the OoC. Previous histological studies have demonstrated that ears with erect OoCs have supporting cells, while flattened or missing OoCs lack supporting cells (22, 26-29). We also visualized large empty spaces within the OoC that likely represent OHC loss (16). Thus, OCT imaging through an intact RWM can estimate the existence or loss of supporting cells, which will be crucial for regenerative therapy.

The present dataset is comprised of three high-integrity specimens (Ears A, B, C) with large OoCs, representing existence of supporting cells. The other two (Ears D, E) have flattened (height < 40 µm) or missing OoCs, and thus lack supporting cells (16, 22, 26-29). Our results suggest that the anatomical integrity of the OoC also impacts how the cochlear partition moves in response to sound.

### Sound-driven motion of tympanic surface of the cochlear partition

#### Motion of the osseous spiral lamina

In the present study, our measurements of the OSL reproduce previous reports of human basal OSL vibration in response to sound stimulus (9, 10). In humans, the basal OSL is very thin, long, and porous; these morphological attributes were previously hypothesized to increase the compliance of the OSL allowing for substantial OSL displacement (8, 11, 30). In previous work, LDV measurements of the tympanic surface of the OSL suggested that it moved like a cantilever plate, where the modiolar end of the OSL was anchored and the distal end (at the OSL-Bridge-J) moved the most (10).

Our results suggest that the displacement magnitude of the OSL at BFpassive differs across ears depending on OoC anatomical integrity (Fig. 5). As can be observed with OCT imaging, cochlear partitions with poor anatomical integrity (with flattened OoC or no OoC, Ears D and E, as seen in Fig. 4) exhibit large OSL displacements (∼22 dB re stapes displacement). Cochlear partitions with high anatomical integrity (with large OoC, Ears A and B) have relatively small OSL displacements (∼12 dB re stapes displacement). The OSL re stapes displacement of Ear C is ∼15 dB and reflects how the integrity of Ear C is in-between that of Ears B and D, as it has a large OoC that appears to have empty space within it (likely due to missing hair cells).

A potential reason for the large OSL motion in ears with low anatomical integrity is that there is a decrease in damping from the OoC, allowing for increased motion of the OSL. Another possibility is that the OSL motion is affected by the mechanical properties and motions of other structures of the cochlear partition. For example, changes in the mass and/or stiffness of connected structures such as the OoC and limbus could affect OSL motion. Ears with poor anatomical integrity could alter the tuned frequency of the BM, thus affecting the motion of the whole cochlear partition—including the OSL. Previous modelling and experimental work has explored such a possibility in gerbils (31, 32). Their results suggested that a loss of cells in the OoC may impact both the stiffness and mass of the cochlear partition, shifting the BF as a result. Such a change may provide an explanation for the differences in motion between the high integrity and low integrity ears in the present study.

Previous LDV measurements of human OSL motion have been reported in 2003 by Stenfelt et al. and 2019 by Raufer et al. (9, 10). In both studies, the anatomy of the OoC was not visualized and thus anatomical integrity was unknown. The large OSL motions reported by Stenfelt et al. are consistent with data from ears with low anatomical integrity in the current study (9). Given that the OSL motion measurements in Stenfelt et al. (9) were from 4-6 days postmortem specimens where the otic capsule was drilled open, it is likely that the OoCs in their specimens were deteriorated and not intact. The specimens in Raufer et al.’s study were 2-4 days postmortem and exhibited a wide variation of OSL motions, similar to our present study (10). In retrospect, it seems likely that the differences in OSL motion seen by Raufer et al. (10) may also have been due to varied OoC anatomical integrity.

#### Motion of the basilar membrane

OCT vibrometry through the intact RWM shows that the cochlear partitions’ BFpassive near the RWM region are between 11-13 kHz. Although this is a relatively narrow range in frequency, this BFpassive variation may be due to differences in the longitudinal location (e.g. a <1.3 mm difference could result in a variation in BFactive from 11-13 kHz), differences in cochlear size, or differences in anatomical integrity across ears.

We find that the radial profile’s point of maximum BM displacement is ∼25 µm distal to the Bridge-BM-J, near the foot of the outer pillar cell (Figs. 5-7). Previous studies that did not have imaging feedback roughly estimated the maximum motion to be at the Bridge-BM-J (10), but the present study more accurately measures it near the foot of the outer pillar cell. The BM and bridge contain continuous, radially-projecting collagen fibers, whose distal and modiolar ends are respectively anchored at the spiral ligament and the OSL (11). Although the bridge and BM contain continuous collagen fibers, the location of maximum motion is not at the center of their combined widths but is instead biased towards the BM. This may be because the limbus attaches to the modiolar aspect of the bridge, stiffening that region. Consequently, the entire radial width of the BM and the distal aspect of the bridge move together in a plate-like fashion, with the distal end of the plate located at the spiral ligament. This plate-like motion, combined with the motion of the OSL and medial portion of the bridge, result in maximum motion near the outer pillar-cell foot (e.g. Fig. 6). The outer pillar-cell foot region having maximum motion could affect shear between RL and TM, and consequently hair-cell stereocilia transduction.

Our data indicates that the motion ratio BM/OSL may also be affected by cochlear integrity. To study this, the motion ratios between the points at the tympanic surface of the cochlear partition and the OSL-Bridge-J are plotted for BFpassive (Fig. 5b). In Ear A, with the highest integrity and the largest OoC, the BM/OSL motion ratio is large (∼19 dB), whereas in Ear E, with low integrity and no OoC, the BM/OSL motion ratio is small (∼3 dB). Ears B-D, which have cochlear integrities in-between Ears A and E, exhibit BM/OSL motion ratios between 8-11 dB (in-between the BM/OSL ratios of Ears A and E). However, between Ears B and D, the integrity-size of the OoC and BM/OSL ratios do not correlate. Our data shows that the ear with the largest OoC has a large BM/OSL motion ratio while the ear with no OoC has a small BM/OSL motion ratio.

### Sound-driven motion of cochlear partition structures in the passive cochlea

In the present study, we also measure displacement characteristics of cochlear partition structures beyond the scala tympani surface towards the scala vestibuli, such as the reticular lamina. We find that the relationship between the RL and BM motion appears to depend on the integrity of the OoC. In ears with high integrity (erect OoC, e.g. Fig. 6), the radial profile of the BM displacement exhibits a simple first mode bending plate behavior, as demonstrated by the “ꓵ”-shape of the magnitude and a flat and smooth phase. In ears with high cochlear integrity, the RL motion is smaller in magnitude, and instead of simple smooth plate-like motion, it has small fluctuations in its magnitude and phase with respect to radial location. In these ears with high cochlear integrity, the RL displacement is also about 5-10 dB less than the BM displacement near BF (Fig. 6 & 7, Ears A-C).

Our findings appear consistent with the passive cochlear partition motion being dominated by BM motion. The BM contains radially projecting collagen fibers, resulting in the plate-like motion as seen in the radial profile. The RL is driven by the BM motion, resulting in lower RL motion than the BM. This observation of the motion of RL being lower than BM has been demonstrated in passive cochlear models of mouse with simplified anatomical details (33). Differences in BM and RL motion could be due to damping from the OoC structures and/or fluid between the BM and RL or alternatively, by differences in BM and RL stiffness. The BM radial collagen fibers move like a beam where stiffness is dominated by beam bending. The RL on the other hand is composed of a mosaic of OHC cuticular plates and tops of phalangeal processes held together by tight junctions. The RL was recently hypothesized to bend and stretch, and thus its stiffness could be dominated by stretching stiffness (18). The RL stretching stiffness is possibly greater than the BM bending stiffness, resulting in less RL motion than BM motion and more complexity in the phase of the RL than the BM. Another possibility for the differences in transverse motion between the BM and RL in the passive case may be related to shearing between the BM and RL instead of compression.

In ears with a flattened or no OoC as in pathological ears (Fig. 7 Ears D and E), the RL and BM have similar displacements and move together. This is likely because the RL and BM are close together, resulting in greater coupling between them.

When the TM is close and likely attached to the RL, as seen in cases with high integrity OoC, the TM and RL have similar displacements (Figs 6&7, Ears A-C). However, in ears with low anatomical integrity (Ears D and E), the TM is not close to or attached to the RL. In these ears, the TM motion is ∼10 dB lower than the RL (as in Fig 7, Ears D and E). In this case, the TM is driven by the local fluid pressure created by the RL. Thus, vibrometry data appears to provide further evidence of functional differences between anatomically healthy and pathological ears.

### Area changes in the human OoC

The observation that the RL motion is less than the BM motion in ears with high cochlear integrity (Ears A and B) indicates that there are changes in the OoC cross-sectional area as it vibrates. Due to the incompressibility of fluid, such an area change would indicate longitudinal fluid flow in the space between the RL and BM. Such longitudinal fluid flow could take place within the cortilymph within the tunnel of Corti, spaces of Nuel, and the outer tunnel.

Our observation of radial cross-sectional area changes in the OoC in human agrees with earlier predictions of longitudinal fluid flow. Guinan et al. (34) previously hypothesized that cyclic OHC motile forces generate OoC area changes, resulting in longitudinal fluid motion. This motion was hypothesized to contribute to amplify the traveling wave near BFactive (see also (35)). To understand how narrow-band tuning can arise from wide-band OHC electromotility, Guinan et al. (34) adapted a simple lumped-element “toy” model of OHC-induced OoC deformation developed by Altoé et al. (36). This adapted toy model incorporated an impedance that is experienced by OHCs when fluid is forced along the narrow longitudinal fluid tunnels (such as tunnel of Corti, spaces of Nuel, and/or outer tunnel). Importantly, this impedance varies with the wavelength of the traveling wave, and the impedance is small only near BFactive where the wavelength is short. As a result, substantial OoC area changes and RL motions occur near BFactive, producing narrow-band traveling-wave amplification (34). Although our human data is passive and thus without OHC electromotility, the concept of longitudinal fluid flow within the OoC is indirectly observed.

### Comparison of OoC measurements in animals and human

The mechanics of the dead passive cochlea differ substantially from the live active cochlea. As opposed to the active cochlea, which is sensitive and sharply tuned to BFactive, the passive cochlea is broadly tuned and exhibits maximal motion at BFpassive that is approximately ½-octave below BFactive (37). Because the measurements in the present study are from passive human cadaveric ears, we compare previous *in vivo* animal measurements at frequencies ½-octave below BFactive, that is at BF-1/2 Oct, where passive mechanics dominate in a live animal.

The OSL is considered to be stationary in the classic view of cochlear mechanics, which was informed by basal measurements from various laboratory animals (e.g. mouse, gerbil, guinea pig, chinchilla, cat). However, there were reports of OSL motion in species such as the squirrel monkey and chinchilla for frequencies below BFactive and not at the round-window region, but rather the more apical region with BFactive ∼9 kHz (4, 38).

To compare the human data in the present study to past animal data, we focus on the live animal’s ratio of motion BM/OSL at BF-1/2 Oct (the frequency ½ octave below BFactive). For example, in the live chinchilla, the BM/OSL ratio of motion was reported to be 20-40 dB at BFactive (9 kHz region), but only 5-15 dB at lower frequencies near live BF-1/2 Oct (38). The BM/OSL ratio at BF-1/2 Oct of previous live animal studies are comparable to the human passive (at BFpassive) BM/OSL ratio of 8-19 dB we observe in Ears A-C, which have good anatomical integrity.

Relative motions of the BM and RL can also be compared across postmortem animal and human data by analyzing the BM/RL magnitude ratio at postmortem BFpassive (frequency of postmortem maximum motion of BM). At postmortem BFpassive, a BM/RL ratio of approximately 2-5 dB was measured from the gerbil basal region less than 60 minutes postmortem (18). In human cadavers we measure BM/RL motion ratios between 3-10 dB in the three ears with high OoC integrity (Ears A-C, Figs. 6-7), and ratios closer to 0 dB in ears with low OoC integrity (Ears D-E, Fig. 7). In summary, despite the use of cadaveric human ears with postmortem times of more than 12 hours, ears with high OoC integrity have OoC motions that are similar or larger compared to postmortem laboratory animals minutes after death.

Consistent with earlier observations of the radial profile of motion, we find the postmortem human maximum motion of the BM to be near the location of the outer pillar cell foot. Other studies in gerbils and guinea pigs have also estimated the point of maximum motion within the BM to be at the outer pillar cell foot or first row of Deiter’s cells. This location of maximum motion was similar both in live and postmortem animals (1, 39-41). It is interesting that there is consistency in the maximum location of BM motion across these species, despite the differences in the anatomy and cochlear partition motion between them. Human specimens have large motions of the OSL and bridge at the base, whereas the gerbil and guinea pig OSL do not move substantially (1, 10, 39-41). Furthermore, the basal cochlea of the gerbil and guinea pig lack a bridge region (10).

### Speculation about OSL and bridge motion affecting the auditory nerve and tuning

The OSL consists of two parallel thin bony plates bridged by a delicate and cavernous network of bony pillars (11, 30). Through this network between the OSL plates, the type 1 auditory nerve fibers (**ANFs**) course radially, projecting from the spiral ganglion to the inner hair cells of the OoC. Near the inner pillar cell feet, the ANFs converge as bundles and pass through small channels located at the vestibular surface of the bridge, the habenula perforata. As they pass through the habenula perforata, type 1 ANFs transition from being myelinated to unmyelinated. Because the region around the habenula perforata moves in human, the question arises whether ANFs are potentially vulnerable at this location. Since it has been demonstrated that small displacements at this location can alter the response properties of motion-sensitive ANF (42), is it plausible that large motions of unmyelinated ANFs could lead to synaptopathy over an extended period, thus related to aging or noise damage?

The relationship between motion of the human cochlear partition structures and ANF response can be further explored in modeling studies. Previously, in a finite-element model, we showed that as ANFs pass between the OSL plates, potential strain and resultant mechanically-induced effects on ANF activity are minimal as they pass between the OSL plates (20). This is because the fibers are located within the neutral axis between the OSL plates, thus experiencing minimal strain. However, the potential effect of motion near the habenula perforata on unmyelinated ANF activity is unknown. The increased OSL motion due to low-integrity OoC may accentuate this problem.

Another interesting observation is that previous psychophysical experiments suggested that cochlear tuning is sharper in humans than in other laboratory animals (43, 44). A potential explanation for this may lie in the anatomical differences between human and laboratory animal cochlea; the basal human cochlea has a vibrating OSL and bridge, as opposed to the base of laboratory animals where the OSL does not vibrate and there is no bridge region (Supplemental information of (10)). As a result, the “effective BM width” may be considered wider in humans due to the motion of the OSL and the bridge. This is consequential, as previous analytical work has predicted that the width of the basilar membrane is related to the spatial spread of tonal signal power (45) and summarized in (46)). This implies that in human the wider effective BM might contribute to sharper mechanical tuning. Such hypotheses can be investigated with computational models that simulate details of human anatomy and motion. Future work of coupling experimental imaging and vibrometry with computational modeling will hopefully answer these questions. Understanding how different human cochlear structures move relative to each other in the passive state will enhance our comprehension of the fundamental mechanics of the human cochlea and the auditory nerve, and how they are affected with pathology.

### Summary

We can visualize and measure vibrometry using OCT to examine the structures of the cochlear partition in intact human cadaveric cochlear specimens without causing damage through the intact round window membrane. OCT imaging can also be used to determine variations in the anatomical integrity of the OoC. Furthermore, cochlear partition motion appears to be linked to the integrity of the OoC. Ears with higher OoC integrity exhibit less vibrations of the OSL around BFpassive. Analysis of BM/OSL vibration ratio shows that the highest integrity (largest OoC) has the largest BM/OSL vibration ratio. This high BM/OSL ratio is similar to findings in previous animal studies. Of clinical importance, minimally invasive determination of the anatomical integrity of the OoC with OCT imaging may enable estimations of supporting cell existence, an important criterion for the success of targeted regenerative therapies. Also, our present study suggests that we can estimate some mechanical function from OoC anatomical integrity. With future technical advancements, perhaps clinical OCT imaging and vibrometry may be feasible.

## Acknowledgements

Supported by grants R01DC013303, F31DC022795, T32DC000038 (Training for Speech and Hearing Sciences), and T32DC000020 (Research Training for Otolaryngology) from the NIDCD/NIH, and the Amelia Peabody Scholars Fund.

Preliminary and partial results in this manuscript were reported in a conference proceedings (47).

## Data Availability

Data from this article will be available on the Harvard Dataverse site, or from the corresponding author on request.

## Author Contributions

P.S., C.I.M., S.P, and H.H.N. conceived of and designed the project. Methodology was developed by C.I.M, N.H.C., and H.H.N. Data was collected by P.S., N.H.C, C.I.M., and H.H.N. Data analysis was performed by P.S., with initial processing and analysis code developed by A.Z. The initial draft of the manuscript was prepared by P.S., S.P., and H.H.N, with feedback from discussions with A.T. and A.Z. All authors reviewed and edited the manuscript.

